# Successor representation supports structural learning of syllable sequences

**DOI:** 10.64898/2026.10.03.756368

**Authors:** Jiali Liu, Yixiang Wang, Jiayu Feng, Haichao Zhao, Qinghua He, Xuefeng Yang, Baishen Liang

**Author notes:** Corresponding authors: Xuefeng Yang, Baishen Liang. These authors contributed equally to this work (Jiali Liu, Yixiang Wang, Jiayu Feng).

## Abstract

Humans efficiently learn the temporal structure of speech, yet the underlying cognitive mechanisms remain unclear. Recent research in visuospatial sequential memory has proposed the successor representation (SR), important in reinforcement learning, which encodes multi-step transitional relationships. To test whether SR also supports the structural learning of rapidly changing speech syllable sequences, we adapted a pyramid-structured sequential learning paradigm using syllable streams presented at a natural speech rate (∼ 4 Hz). Participants exhibited robust structural learning, evidenced by asymmetric memory performance modulated by adjacent transitional probabilities. Notably, the SR model captured recognition performance for non-adjacent syllable pairs. Moreover, SR-based learning effects generalized across all syllable sequences, regardless of adjacency. Furthermore, the effectiveness of SR learning was associated with cognitive flexibility and working memory capacity. Together, these findings suggest that humans rely on effortful SR-like mechanisms for the structural learning of syllable sequences, highlighting a modality-general computational principle for sequential learning across various temporal modulation scales.

## 1. Introduction

### 1.1. Syllable sequences in speech

Speech is an essential communication tool in social interactions. While speech embeds linguistic information that is hierarchically organized, it reaches listeners as an almost gapless syllable stream. Therefore, speech comprehension requires the listeners to quickly and effectively segment syllable sequences to reconstruct the constituted linguistic units (Christiansen & Chater, 2016). To grasp this skill for understanding a specific language, humans need to acquire not only its syllable system, but also the temporal transitions between syllables (e.g., /LO/ is expected when /HEL/ is heard).

The simplest explanation for the acquisition of syllable transitions is the Hebbian law, where the “fired together, wired together” rule implies that consecutive syllables are organized into an integrated unit stored in memory (Hebb, 1961). However, natural language is constituted by multiple layers of non-adjacent dependencies (NADs) (Petkov & Ten Cate, 2020), including temporal structures of syllables (e.g. the “is” always predicts “ing” regardless of the intervening elements, as in “the bird is singing”). Previous studies have found evidence supporting the learning of NADs in speech (Endress & Bonatti, 2007; Wang et al., 2019). However, these studies are usually short and apply rigid and deterministic dependency frames (e.g., A-X-B, where A and B co-occur, but the X is highly variable), which might have induced rule-based learning that weakened ecological validity for real-world language processing. Especially, natural languages contain multi-step and probabilistic-based dependencies (e.g., the term fowling “is” can either be a present progressive verb “singing” or an adjective “red”) (Mersad & Nazzi, 2011). Meanwhile, the intervening elements in natural languages can also act as structural nodes for the dependencies (e.g., the “and” in “the bird is singing and jumping” links the “is” and the “-ing” in “jumping”). These considerations call for approaches that can capture multi-step, probabilistic predictive relationships within syllable sequences.

### 1.2. Insights from spatial navigation and cognitive maps

Converging evidence from spatial navigation and event-sequential memory studies indicates that humans possess domain-general structural learning capacities that extend beyond local association, yielding internal relation representations that are often conceptualized as cognitive maps(Behrens et al., 2018). Building on this framework, two broad representational hypotheses have been proposed regarding how relational structure is represented. First, Euclidean map accounts posit that relational structure is embedded in a geometric space (Peer et al., 2021), consistent with evidence that hippocampal–entorhinal regions encode Euclidean distance– and direction-related signals during spatial navigation and in abstract representational spaces (Howard et al., 2014; Liu et al., 2023). By contrast, the successor representation (SR), an important approach in human reinforcement learning (Momennejad et al., 2017), characterizes relational structure in predictive terms, enabling multi-step prediction by representing each state via its expected future occupancy across the environment(Dayan, 1993; Gershman, 2018).

These proposals are highly relevant to speech by supporting inference over non-adjacent relations and multi-step structure. However, a critical gap is that most evidence for SR-like or map-like representation comes from relatively slow-paced visual/spatial tasks(Tacikowski et al., 2024), whereas speech syllables fluctuate at ∼4–5 Hz. Constructing and exploiting SR- or map-like representations may be cognitively demanding, raising the possibility that such model-based computations are difficult to deploy online under stringent temporal pressure (Otto et al., 2015; Seel, 2017). Accordingly, it remains unresolved whether inference in continuous syllable streams is better characterized by predictive coding (consistent with SR), geometric distance coding (consistent with Euclidean cognitive maps), or a fallback to heuristic associative chaining.

### 1.3. The current study

In the present study, we investigated how humans learn and represent the temporal structure of syllable sequences in continuous speech-like input. By adapting a pyramid-structured sequential learning paradigm originally developed in the visual domain (Tacikowski et al., 2024) to ∼4 Hz syllables streams, we examined whether structural representations proposed in slower-paced domains can be formed and exploited under the temporal constraints of speech processing. To determine whether the structural representation had emerged, we analyzed adjacent syllables and hypothesized that if participants engaged in structural learning rather than simple chunking, their response accuracy would be asymmetric for inner-outer versus outer-inner pairs (**Figure 3a**). To further characterize the specific nature of this structure, we then examined non-adjacent syllables, testing whether performance was better predicted by SR (predictive coding) or Euclidean distance (geometric coding) (**Figure 4-5**).

**Figure 1.**
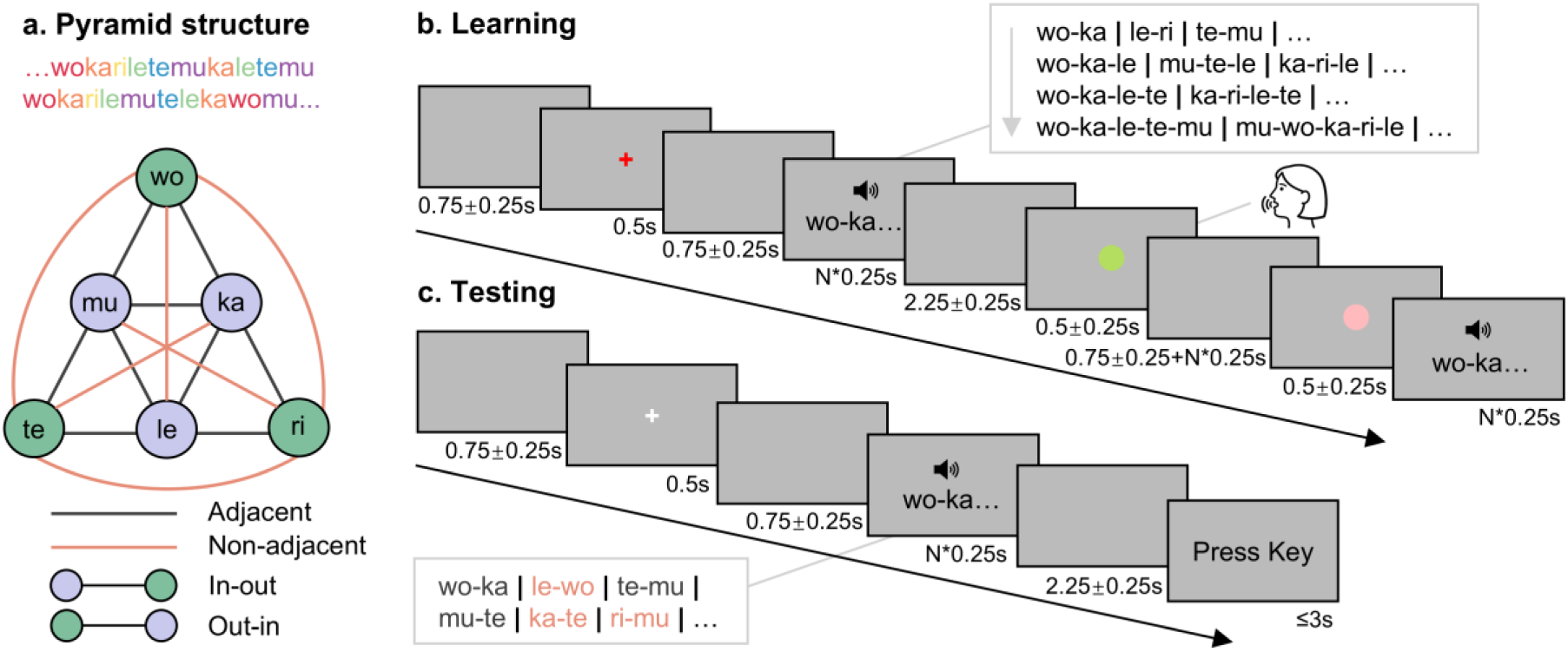
Stimuli and task. (a) The temporal sequences of the syllables (top) and the pyramidal structure for temporal transitions. Purple circles represent inner syllables, which have four adjacent connections; green circles represent outer syllables, which have only two adjacent connections. Black lines indicate adjacent transitions; coral-colored lines indicate non-adjacent transitions. (b) Learning phase. Participants learned syllable sequences composed of consecutive 2, 3, 4, and 5 syllables, where N denotes the number of syllables in the sequence. (c) Testing phase. Participants were required to recognize whether syllable pairs had appeared during the learning phase.

**Figure 2.**
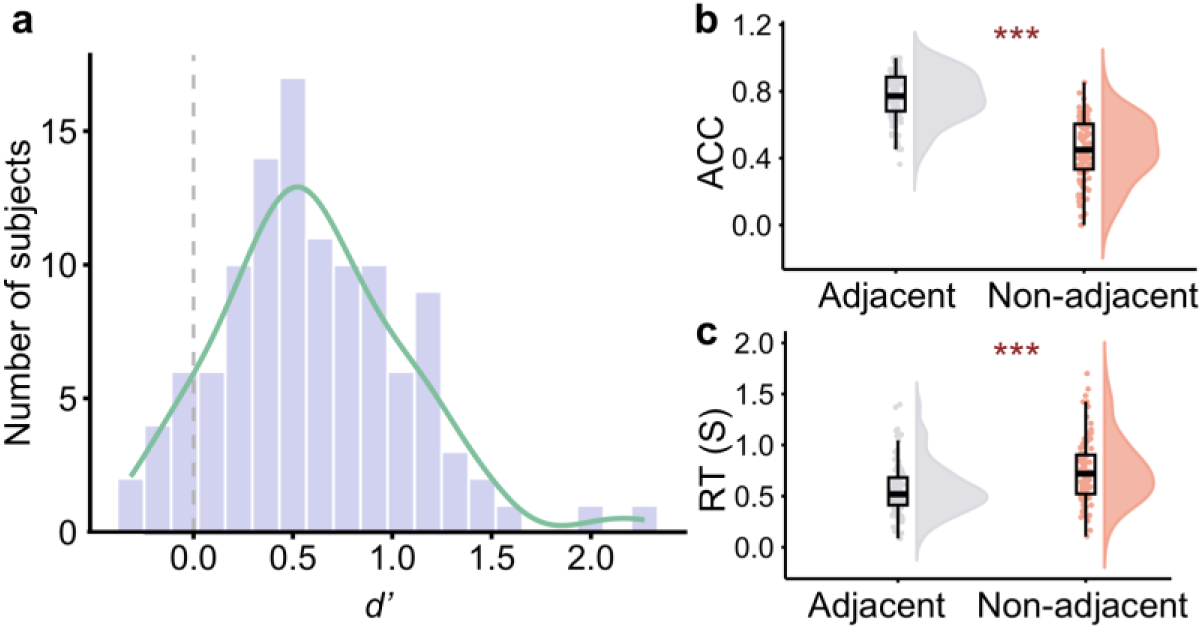
Task performance. **(a)** Distribution of sensitivity (*d*’) in the task. The distribution indicates that most participants successfully discriminated the learned consecutive syllable pairs from the unlearned distractors. **(b)** Mean accuracy (ACC) and **(c)** mean reaction time (RT) for adjacent and non-adjacent pairs. Participants showed significantly higher accuracy and faster responses (shorter RTs) for adjacent pairs compared with non-adjacent pairs. Boxplots show the median (center line), the interquartile range (IQR; 25^th^–75^th^ percentiles), and the minimum and maximum values within 1.5 × IQR. * *p* < .05, ** *p* < .01, *** *p* < .001.

**Figure 3.**
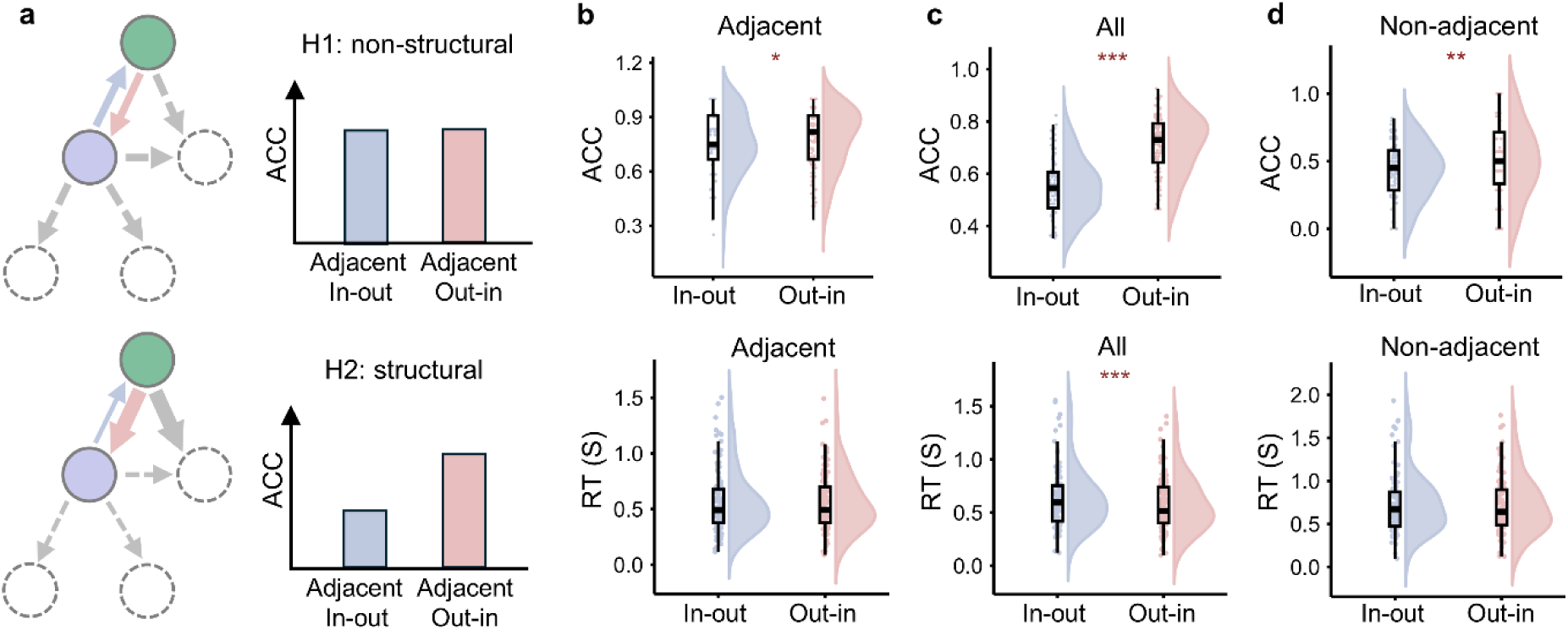
Structural learning of syllable sequences. **(a)** Schematic illustration of the hypotheses. The non-structural learning hypothesis (Hypothesis 1) predicts equal performance (upper), whereas the structural learning hypothesis (Hypothesis 2) predicts lower accuracy for in-out pairs (lower). Light blue and light pink arrows represent in-out and out-in pairs, respectively. Arrow thickness indicates transitional probability, which is hypothesized to be associated with ACC (thin: low TP; thick: high TP). **(b-d)** Mean accuracy (ACC) and reaction time (RT) for in-out and out-in pairs in the **(b)** adjacent, **(c)** all-pairs, and **(d)** non-adjacent conditions. Across all conditions, in-out pairs yielded significantly lower accuracy pairs relative to out-in pairs. In-out pairs also exhibited significantly longer RT than out-in pairs for all-pairs condition. Boxplots show the median (center line), the interquartile range (25^th^–75^th^ percentiles), and the minimum and maximum values within 1.5 × IQR. * *p* < .05, ** *p* < .01, *** *p* < .001.

**Figure 4.**
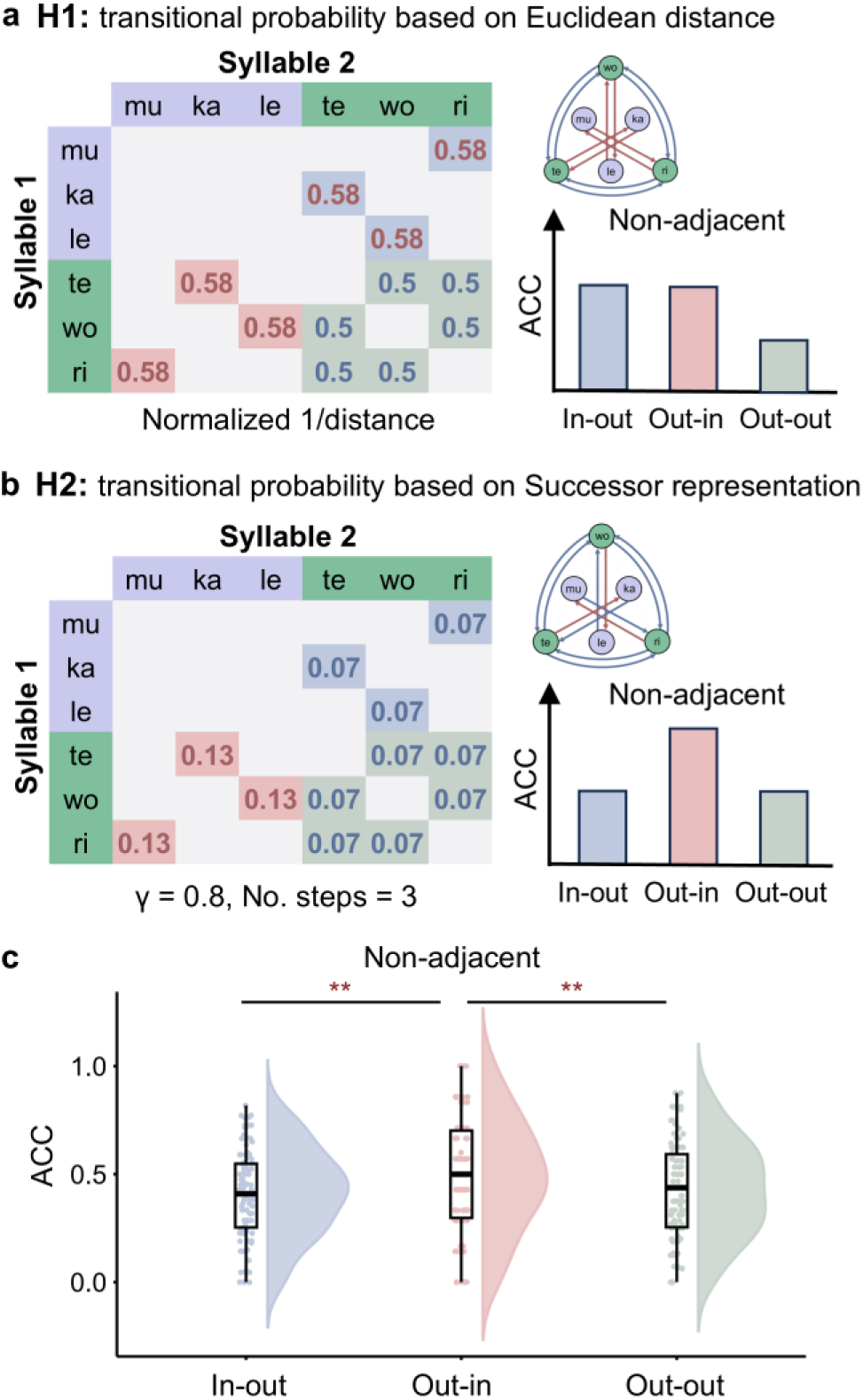
Successor representation (SR), but not geometric representation (GR), supports structural learning. **(a)** Hypothesis 1: GR model. Under this hypothesis, sequence structure is represented geometrically, such that transitional probabilities are defined by the normalized inverse Euclidean distance. This model predicts equivalent performance for out-in and in-out pairs, both higher than out-out pairs for the non-adjacent condition. **(b)** Hypothesis 2: SR model (γ= 0.8, steps = 3), in which transitional probabilities reflect predictive relationships between states. The SR model predicts that higher performance for out-in pairs relative to in-out and out-out pairs, which should not differ, in the non-adjacent condition. Cell shading represents pair types (light blue: in–out; light pink: out–in; light green: out–out). Text color indicates transitional probabilities (dark pink: high; dark blue: low). Purple and green headers denote inner and outer syllables, respectively. **(c)** ACC for non-adjacent pairs. Accuracy was significantly higher for out-in pairs compared with both out-out and in-out pairs; the latter two did not differ significantly, which is exactly consistent with the predictions of the SR model. Boxplots show the median (center line), the interquartile range (25^th^–75^th^ percentiles), and the minimum and maximum values within 1.5 × IQR. * *p* < .05, ** *p* < .01, *** *p* < .001.

**Figure 5.**
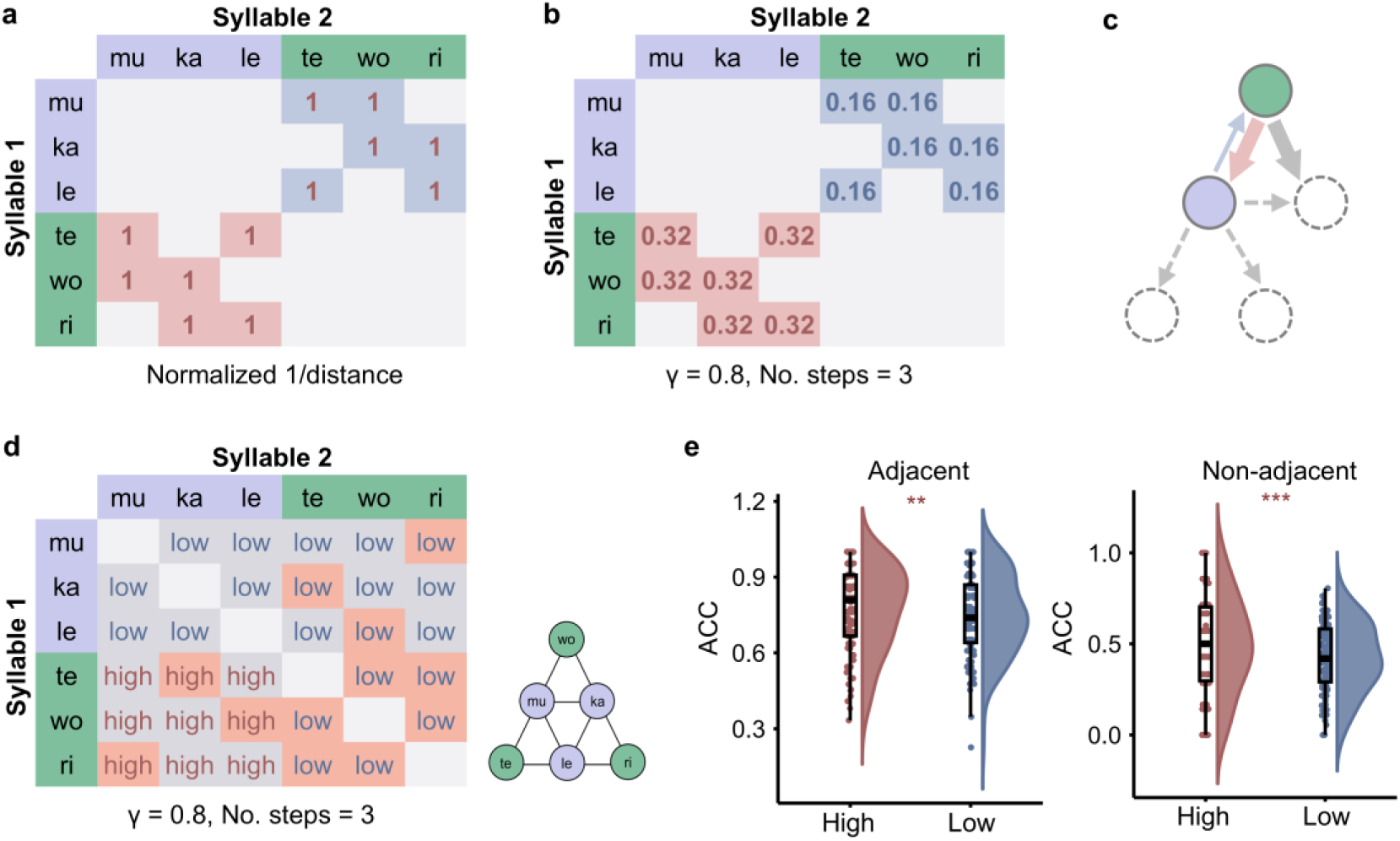
Unified successor representation mechanisms support the structural learning of adjacent and non-adjacent syllable sequences. **(a-b)** Transitional probabilities (TPs) for adjacent pairs derived from the **(a)** GR model and **(b)** SR model (γ = 0.8, steps = 3), respectively. Light blue and light pink shading represent in–out and out–in pairs. Dark pink and dark blue text indicate high and low TPs. **(c)** Schematic of the SR architecture for adjacent pairs, where thick arrows represent high TPs, and thin arrows denote low TPs, and arrow colors indicate pair types (light blue: in–out, light pink: out–in). (**d**) Matrix of TPs within the SR space, where TP values were dichotomized into low and high groups for both adjacent and non-adjacent pairs. Gray shading represents adjacent transitions; coral shading represents non-adjacent transitions. (**e**) Accuracy (ACC) for trial pairs with different TPs. For both adjacent and non-adjacent conditions, pairs with high TPs showed significantly higher accuracy than those with low TPs. Boxplots show the median (center line), the interquartile range (25^th^–75^th^ percentiles), and the minimum and maximum values within 1.5 × IQR. * *p* < .05, ** *p* < .01, *** *p* < .001.

## 2. Methods

### 2.1. Participants

A total of 115 right-handed participants (19 males, 96 females; mean age = 20.63 years, SD = 1.84) were recruited for this study. All participants reported normal or corrected-to-normal vision and normal hearing, with no history of color blindness or color weakness. They reported no history of neurological or psychiatric disorders and were not taking any medication at the time of the experiment. All participants provided written informed consent and were compensated for their participation. The study was approved by the Ethics Committee of Southwest University.

### 2.2. Materials

The learning materials consisted of six phonetically grammatical but meaningless Mandarin syllables: [tə^55^], [kɑ^35^], [ʐɭ^55^], [lə^35^], [mu^55^], and [wɔ^35^], corresponding to the Pinyin tē, ká, rī, lé, mū, and wó. These syllables were recorded by a 26-year-old female native Mandarin speaker in a quiet room, using a UM2 USB external sound card with a sampling rate of 44,100 Hz. Audio files for the stimuli were selected to ensure consistent volume, envelope shapes, and sound quality. Audacity software was used to adjust pitch height and contour for the high-level [55] and rising tones [35]. Each sound file was initially 500 ms long and then compressed to 250 ms while maintaining fidelity. Root-mean-square (RMS) balancing in Praat software ensured consistent energy levels across stimuli.

The selected Mandarin syllables and their combinations had the following linguistic characteristics: (1) the consonants covered the common manners of articulation in Mandarin, including plosives (e.g., [t], [k]), fricatives (e.g., [ʐ]), laterals (e.g., [l]), and nasals (e.g., [m], [w]), (2) the vowels included main Mandarin vowels (i.e., [ə], [ɭ], [u], [ɑ], and [ɔ]), (3) all syllables were meaningless and did not correspond to any existing Chinese characters; and (4) consecutive syllable pairs did not form meaningful Mandarin words in the learning phase. These guarantee that the stimuli contain sub-syllabic structures resembling natural languages, while keeping syllables meaningless.

Syllable presentation followed a pyramidal structure (see **Figure 1a**). After a syllable was presented, only those directly connected to it by connecting links could appear next. For example, "wo" could follow "mu" (black line in **Figure 1a**), but not "le" (red line). Participants were instructed to learn this transitional temporal structure between syllables. We chose this pyramidal structure for several reasons. First, it has been verified by previous studies in visual sequential learning and helps build direct comparisons. Second, it encapsulates the probabilistic-based transitional relationships of both adjacent and non-adjacent syllables in natural languages, instead of the strictly rule-based dependencies. Third, the nested structure with both inner and outer nodes parameterize inter-syllable transitional probabilities (TPs), ensuring direct estimation of TPs dissociating from syllables. Nevertheless, syllables were presented at a transitional (modulation) rate of 4 Hz (i.e., 4 syllables per second), substantially faster than the past sequential learning paradigms, to align with natural speech (Poeppel & Assaneo, 2020).

### 2.3. Procedure

The experiment consisted of a learning phase and a testing phase, both presented using Psychtoolbox. To control for circadian effects, all participants completed the learning phase at 10:00 am. A recognition test was given ten minutes after the learning phase to assess immediate memory performance.

#### Learning Phase (Figure 1b)

Participants learned syllable sequences consisting of two (18 combinations, e.g., wo-ka), three (42 combinations, e.g., wo-ka-le), four (66 combinations, e.g., wo-ka-le-te), and five (66 combinations, e.g., wo-ka-le-te-mu) syllables. Each combination was presented in both forward and reverse orders (e.g., "wo-ka" and "ka-wo"), resulting in 192 trials, repeated twice for a total of 384 trials.

In each trial, a red “+” symbol appeared in the center of the screen for 0.5 s to ensure participants’ attention. The syllable sequence was then presented, with each syllable lasting 0.25 s (e.g., a two-syllable combination lasted 0.5 s, a three-syllable combination lasted 0.75 s). The syllables were presented audibly, with no text displayed. Following the auditory presentation, a green circle was represented to prompt participants to orally repeat and memorize the syllable sequence. Simultaneously, a red circle appeared with the playback of the correct syllable sequence, serving as a reference for participants to verify their repetition accuracy. All stimuli were displayed on a gray background. To reduce anticipation effects, a random jitter of ±250 ms was added to the timing of blank screens and stimuli (see **Figure 1b** for detailed timing). Participants were not explicitly informed of the pyramidal structure of the sequences.

#### Testing Phase (Figure 1c)

Participants completed a recognition task in which they judged whether the presented syllable pair had appeared during the learning phase. The syllable pairs could be those presented during learning (adjacent combinations appeared during the learning tasks, e.g., "wo" and "ka," indicated by black lines in **Figure 1a**) or not presented (non-adjacent combinations, e.g., "wo" and "le," indicated by red lines in **Figure 1a**). Both types were presented equally for a total of 90 trials. Participants were instructed to respond as quickly and accurately as possible by pressing the ‘F’ key if they recognized the combination and the ‘J’ key if they did not. If the reaction time exceeded 3 seconds, the trial was automatically skipped and recorded as an error. Practice sessions were conducted before both the learning and testing phases to ensure participants fully understood the procedure and key operations.

#### Follow-up Assessment

To investigate the potential influence of individual differences on SR performance, we conducted a follow-up session assessing cognitive flexibility and working memory. Cognitive flexibility was measured using the Cognitive Flexibility Inventory (CFI; Dennis & Vander Wal, 2010), a self-report questionnaire evaluating the ability to perceive difficult situations as controllable and generate multiple alternative solutions. Working memory capacity was assessed using the 2-back task, a classic paradigm requiring participants to monitor a series of stimuli and respond when the current stimulus matches the one presented two steps earlier (Jaeggi et al., 2010). Due to participant attrition during the recall process, sample sizes varied across measures: the online administration of the CFI yielded data from 95 participants, whereas the requirement for on-site laboratory attendance for the 2-back task resulted in a final sample of 86.

### 2.4. Successor representation model

The successor representation (SR) model conceptualizes the transitional probability between syllables A and B not as a single step, but as the discounted sum of probabilities across all possible pathways connecting them (Dayan, 1993; Gershman, 2018). For instance, given a finite horizon ***T*** of 3, the possible trajectories from "wo" to "mu" in our experimental design include: (1) the direct path "wo" → "mu", (2) the two-step path "wo" → "ka" → "mu", and (3) the three-step path "wo" → "ka" → "le" → "mu". Crucially, the contribution of each pathway to the overall SR value decays with path length, scaled by a discount factor **γ**.

Here, we adopted a normalized finite-horizon version of SR model. The SR transitional probability matrix among syllable pairs in this study, denoted as ***M_SR_***, was computed as the normalized sum of discounted transitional probabilities by a finite horizon ***T*** (**Formular 1**). In the formular, ***P*** is the one-step transitional probability matrix defined by the reciprocal of the count of possible successor syllables (i.e., 0.25 for pairs starting with inner nodes, and 0.5 for pairs starting with outer nodes, **Table 1** and **Figure 1a**). Meanwhile, **γ** is the discount factor (0 < **γ** < 1). The SR matrix is defined as:

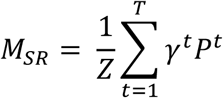

**Table 1.** Consecutive transitional probabilities (finite horizon T = 1)

|  | **mu** | **ka** | **le** | **te** | **wo** | **ri** |
| --- | --- | --- | --- | --- | --- | --- |
| **mu** | 0.0 | 0.25 | 0.25 | 0.25 | 0.25 | 0.0 |
| **ka** | 0.25 | 0.0 | 0.25 | 0.0 | 0.25 | 0.25 |
| **le** | 0.25 | 0.25 | 0.0 | 0.25 | 0.0 | 0.25 |
| **te** | 0.5 | 0.0 | 0.5 | 0.0 | 0.0 | 0.0 |
| **wo** | 0.5 | 0.5 | 0.0 | 0.0 | 0.0 | 0.0 |
| **ri** | 0.0 | 0.5 | 0.5 | 0.0 | 0.0 | 0.0 |

Note. Each cell value represents the transitional probability from the syllable in the row to the syllable in the column.

where ***P^t^*** represents the t-step transitional probability matrix, and 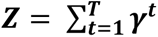 is the normalization constant. In the current study, we set **γ** = 0.8, and ***T*** = 3, matching one step more than the medium path length embedded in the experimental design (max length = 4) while ensuring that more distant transitions contributed less to the SR value. The transitional probability matrix derives from the SR model is shown in Table 2.

**Table 2.** SR transitional probabilities (finite horizon T = 3, γ = 0.8)

|  | **mu** | **ka** | **le** | **te** | **wo** | **ri** |
| --- | --- | --- | --- | --- | --- | --- |
| **mu** | 0.16 | 0.23 | 0.23 | 0.16 | 0.16 | 0.07 |
| **ka** | 0.23 | 0.16 | 0.23 | 0.07 | 0.16 | 0.16 |
| **le** | 0.23 | 0.23 | 0.16 | 0.16 | 0.07 | 0.16 |
| **te** | 0.32 | 0.13 | 0.32 | 0.1 | 0.07 | 0.07 |
| **wo** | 0.32 | 0.32 | 0.13 | 0.07 | 0.1 | 0.07 |
| **ri** | 0.13 | 0.32 | 0.32 | 0.07 | 0.07 | 0.1 |

Note. Each cell value represents the transitional probability from the syllable in the row to the syllable in the column.

Unlike standard transitional probability measures based solely on immediate successors, the SR model captures multi-step predictive relationships, thereby offering a unified quantification for both adjacent and non-adjacent dependencies. However, SR probabilities inevitably exhibit collinearity with adjacency, as adjacent transitions inherently possess higher values than non-adjacent ones. To disentangle the effects of SR probability from adjacency, we employed a stratified binarization approach. Specifically, syllable pairs were classified into ’High’ and ’Low’ SR groups for the Adjacent and Non-adjacent pairs separately. We established specific thresholds to ensure balanced grouping: for non-adjacent pairs, the cut-off was set at 0.10, while for adjacent pairs, the cut-off was set at 0.28 (**Figures 4b and 5b, d**).

### 2.5. Geometric representation model

The geometric representation (GR) model quantifies the transitional relationship between syllables A and B based on their Euclidean distance within the spatial configuration, as depicted in **Figure 1a**. In this framework, the Euclidean distance between adjacent outer nodes (e.g., ’wo’ and ’ri’) was standardized to 1, and distances for all other adjacent and non-adjacent pairs were derived geometrically (i.e., via the Pythagorean theorem). To convert these spatial distances into a metric comparable to transitional probabilities, we computed the reciprocal of the distance matrix (i.e., 1/distance). Diagonal elements (self-transitions) resulting in infinity were reset to zero. Finally, the values were normalized by dividing by the maximum value in the matrix, ensuring a scaled representation where shorter distances correspond to higher transition strengths (**Table 3**).

**Table 3.** Geometric transitional probabilities

|  | **mu** | **ka** | **le** | **te** | **wo** | **ri** |
| --- | --- | --- | --- | --- | --- | --- |
| **mu** | 0.0 | 1.0 | 1.0 | 1.0 | 1.0 | 0.58 |
| **ka** | 1.0 | 0.0 | 1.0 | 0.58 | 1.0 | 1.0 |
| **le** | 1.0 | 1.0 | 0.0 | 1.0 | 0.58 | 1.0 |
| **te** | 1.0 | 0.58 | 1.0 | 0.0 | 0.5 | 0.5 |
| **wo** | 1.0 | 1.0 | 0.58 | 0.5 | 0.0 | 0.5 |
| **ri** | 0.58 | 1.0 | 1.0 | 0.5 | 0.5 | 0.0 |

Note. Each cell value represents the transition probability from the syllable in the row to the syllable in the column.

### 2.6 Statistics

Data analyses were performed using R statistical software (Version 4.5.2). Linear and generalized linear mixed-effects models were fitted using the lme4 package, with p-values estimated via the lmerTest package. Before statistical analyses, trials with reaction times (RTs) exceeding ±2.5 standard deviations from each participant’s mean were excluded to reduce the influence of extreme values. This procedure resulted in the exclusion of 2.52% ± 1.26% of RT trials.

For the Trial-level analyses, we fitted Linear Mixed-Effects (LME) models for reaction times (RTs) and Generalized Linear Mixed-Effects (GLME) models with a binomial distribution (logit link function) for response accuracy (ACCs). All models included a random intercept for each participant (1 | subject) to account for inter-individual variability. Four specific models were constructed to address our research questions: (1) Models with Adjacency (Adjacent vs. Non-adjacent) as the fixed effect were used to assess whether participants successfully learned the sequential structure of the syllable streams. (2) Focused analyses on adjacent pairs were performed with Pair Type (In-out vs. Out-in) as the fixed effect to verify the structural learning hypothesis. (3) To distinguish SR from GR models, we analyzed trials in the non-adjacent condition. Models were fitted with Pair Category (out-out, out-in, and in-out pairs) as the fixed effect. Post-hoc pairwise comparisons were conducted using Estimated Marginal Means (EMMs) to test specific contrasts. (4) Comprehensive models where SR Probability (Low vs. High), Adjacency (Adjacent vs. Non-adjacent), and their interaction were entered as fixed effects to assess the global sensitivity to statistical regularities.

For the subject-level analyses, trial-level data were first averaged for each participant within each condition defined by SR probability and Adjacency. These aggregated measures were used to complement the trial-level mixed-effects analyses. We conducted 2 (SR probability: Low vs. High) × 2 (Adjacency: Adjacent vs. Non-adjacent) repeated-measures analyses of variance (ANOVAs) to assess main effects and interactions. Significant interactions were decomposed using post-hoc pairwise comparisons with Holm correction.

Furthermore, to explore the individual difference, we calculated structural learning indices ΔACC (*Accuracy_Hig_*_ℎ_ *_SR_* -*Accuracy_Low SR_*) and ΔRT (*Reaction Time_Hig_*_ℎ_ *_SR_* -*Reaction Time_Low SR_*) based on the aggregated data. Participants were divided into High and Low groups based on their scores on the Cognitive Flexibility Inventory (CFI) and their performance in the 2-back task. The structural learning indices (i.e. Δ ACC and ΔRT) and cognitive groups were compared between groups using independent-samples t-tests with False Discovery Rate (FDR) correction.

One-sample t-tests were used to determine whether performance differed significantly from chance level (0.5) where relevant. The Holm-Bonferroni method was applied to control for multiple comparisons where appropriate.

## 3. Results

### 3.1. General task performance

We first examined general task performance to validate that participants successfully learned the sequential structure of the syllable streams. Overall, participants performed well above chance in the recognition test (*t*(113) = 14.26, *p* < .001, **Figure 1c**). To evaluate individual response bias, *d′* scores were calculated for each participant (**Figure 2a**). Most participants showed positive *d′* values, and a small proportion (6.67%) with negative scores were excluded to ensure data validity(Peng et al., 2025). Recognition accuracy was higher for adjacent than non-adjacent pairs (*β* = −1.44, *SE* = 0.05, *z* = −31.04, *p* < .001, **Figure 2b**). Reaction time analyses showed a corresponding adjacency effect, with faster responses to adjacent pairs (*β* = 0.16, *SE* = 0.01, *t*(5455) = 16.26, *p* < .001, **Figure 2c**). Given these robust differences, subsequent analyses were conducted separately for adjacent and non-adjacent trials.

### 3.2. Structural learning of syllable sequences

General performance analysis (**Figure 2**) indicated that most participants successfully recognized the learned consecutive syllable pairs (i.e., the adjacent syllable combinations) from the unlearned distractors (i.e., non-adjacent pairs). Building on this result, we further address the question of whether the acquisition of the consecutive syllable pairs was a Hebbian pairing process (i.e., model-free, or non-structural), or participants in fact learned the temporal transitional relationships embedded in the syllable pairs, as constrained by the pyramidal structure (**Figure 1a**).

Particularly, we sub-selected adjacent syllable pairs with inner-outer (in-out) or outer-inner (out-in) position relationship to test whether statistical learning exists. This comparison guarantees the estimation of effects from transitional probabilities (TP; during learning, *in-out* pairs have lower TP than *out-in* pairs because an inner syllable connects to two outer and two inner syllables, whereas an outer syllable connects to only two inner syllables, **Figure 3a**), while controlling for confounds from syllable occurrence frequency (in-out and out-in pairs are symmetrical). GLME analyses revealed a significant difference in accuracy between in-out and out-in pairs in the adjacent condition (*β* = −0.18, *SE* = 0.09, *z* = −2.09, *p* = 0.037; **Figure 3b**, upper panel), while the reaction times (LME) showed no significant difference (*β* = 0.01, *SE* = 0.01, *t*(3413) = 0.55, *p* = 0.586; **Figure 3b** lower, panel). The adjacent in-out and out-in pairs appeared with equal frequency during the learning phase (n = 186 per condition), precluding exposure frequency as an alternative explanation for the accuracy differences. To investigate whether the observed performance differences stemmed from participants basing their judgments exclusively on the first syllable (syllable 1) exclusively, we compared the accuracy for adjacent out-in pairs with the adjacent in-in pairs. The result revealed no significant difference between these pairs (*β* = −0.09, *SE* = 0.09, *z* = −1.01, *p* = 0.312). This null effect indicates that participants’ judgments were not based merely on the first syllable; instead, the identity of the second syllable also played a critical role.

We extended the analysis to the non-adjacent pairs to all pairs (i.e., pooling adjacent and non-adjacent pairs) to evaluate the robustness of the in-out/out-in difference. Consistent with the adjacent results, the accuracy results showed a significant effect between in-out and out-in pairs for both non-adjacent (*β* = 0.29, *SE* = 0.09, *z* = 2.98, *p* = 0.003, **Figure 3d**) and all pairs (*β* = 0.44, *SE* = 0.06, *z* = 7.19, *p* < 0.001, **Figure 3c**). For RT, a significant effect was observed only for all pairs (*β* = -0.11, *SE* = 0.01, *t*(5431.22) = -7.229, *p* < 0.001), whereas the non-adjacent pairs did not reached significance (*β* = -0.04, *SE* = 0.02, *t*(1921.67) = -1.634, *p =* 0.102).

### 3.3. Successor, but not geometric representation, supports structural learning

Given that structural learning was found, a further question to be addressed was whether such structural learning is better supported by predictive coding (consistent with successor representation) or by geometric coding (strictly model-based; consistent with Euclidean cognitive maps). To address this question, we sub-selected testing trials with non-adjacent syllable pairs for analysis. Noted that the non-adjacent syllable pairs were not present during the learning phase but, based on the specific pyramidal structure used in our study, could still be subcategorized into three types: outer-outer (out-out), outer-inner (out-in), and inner-outer (in-out) pairs.

Recognition accuracies from the out-in pairs were significantly higher than those from the out-out pairs (GLME pairwise comparisons: *β* = 0.29, *SE* = 0.10, *z* = 3.13, *p* = 0.003, **Figure 4c**), confirming that structural learning can be generalized into the unlearned, non-adjacent pairs during the learning phase. Both the successor representation (SR) and geometric representation (GR) models successfully captured this effect (**Figure 4a, b**). Specifically, the out-in trials were categorized into the high TP group in both SR (TP = 0.13) and GR (TP = 0.58) models, while the out-out pairs were categorized into the low TP group in both SR (TP = 0.07) and GR (TP = 0.5). Therefore, the recognition accuracy for non-adjacent pairs with higher TPs (i.e. out-in) was higher than those with lower TPs (i.e. out-out).

However, the SR and GR models predicted the non-adjacent in-out pairs differently, which derived testable hypotheses linking to the recognition performance. While the GR assigned a high TP for the in-out pairs (TP = 0.58, i.e., Hypothesis 1: in-out = out-in > out-out, **Figure 4a**), the SR assigned a low TP, instead (TP = 0.07, i.e., Hypothesis 2: out-in > out-out = in-out, **Figure 4b**). Pairwise comparisons revealed that the recognition accuracy for the in-out pairs was significantly lower than that for the out-in trials (*β* = -0.31, *SE* = 0.09, *z* = -3.31, *p* < 0.001, **Figure 4c**). Moreover, no significant difference was found between the recognition accuracy for the in-out and the out-out trials (*β* = -0.02, *SE* = 0.07, *z* = -0.25, *p* = 0.806, **Figure 4c**). Collectively, these results support the successor representation model, rather than the geometric representation model. However, the main effect on reaction times by the three types was not significant (*F*(1.80, 203.62) = 0.09, *p* = .890, *η_p_²* = 0.002; **Supplementary Fig. 2a**).

### 3.4. Unified successor representation mechanisms support the structural learning of adjacent and non-adjacent syllable sequences

Given that we found structural learning in the memory of syllable sequences (**Session 3.2**) for adjacent syllable pairs, and that the successor representation, instead of geometric representation, supports structural learning (**Session 3.3**) for non-adjacent pairs, we further asked whether successor representation can provide a unified explanation for structural learning, regardless of adjacency.

We first evaluated whether the successor representation (SR) and geometric representation (GR) models could account for the behavioral asymmetries observed between in-out and out-in adjacent syllable pairs (**Figure 3**). The GR model failed to differentiate between these conditions, assigning an identical geometric transitional probability of 1 to both groups (**Figure 5a**). This limitation arises because probability estimates derived solely from Euclidean distance are inherently symmetric and lack directionality. In contrast, the SR model successfully captured this directional bias, predicting higher transitional probabilities for out-in pairs (TP = 0.32) compared with in-out pairs (TP = 0.16) (**Figure 5b**). This prediction aligned with the behavioral results, where higher recognition accuracy and faster reaction times were observed for syllable pairs associated with higher SR probabilities (i.e., the out-in condition) (**Figure 3**).

We then pooled the adjacent and non-adjacent syllable pairs to investigate whether the SR model could provide a unified account of recognition performance, irrespective of adjacency. To this end, transitional probability matrices were computed for all potential syllable pairs using the SR model and subsequently dichotomized into Low and High groups. (see **Methods, 2.4 & 2.5, Figure 5d**). We then analyzed participants’ response accuracy (ACCs) and reaction times (RTs) as a function of SR transitional probability (Low vs. High) and adjacency (Adjacent vs. Non-adjacent). To ensure the robustness of our findings, we employed both trial-level mixed-effects modeling and subject-level analysis of variance (ANOVA).

For ACCs, the GLME model revealed a significant main effect of SR transitional probability (*β* = 0.14, *SE* = 0.07, *p* = .036, **Figure 5e-f**). Participants showed significantly higher accuracy for pairs with high SR probabilities compared to those with low SR probabilities. We also observed a significant main effect of adjacency, consistent with the finding in **Figure 2** (*β* = -1.42, *SE* = 0.06, *p* < .001). The interaction between SR probability and adjacency was not significant (*p* = .209). For RTs, the LME model yielded a significant main effect of adjacency (*β* = 0.03, *SE* = 0.01, *p*= .002). Participants responded significantly slower to non-adjacent pairs compared to adjacent pairs. However, the main effect of SR probability was not statistically significant (*β* = -0.02, *SE* = 0.01, *p* = .129), nor was the interaction effect (*p* = .290). Crucially, the significant main effect of SR probability, combined with the absence of an interaction with adjacency, indicates that the SR model provides a unified account of recognition performance irrespective of adjacency.

The ANOVA on ACCs revealed a significant main effect of SR probability (*F*(1, 113) = 15.63, *p* < .001, *η^2^* = .014), and a significant main effect of adjacency (*F*(1, 113) = 106.87, *p* < .001, *η^2^* = .378. Different from the LME analysis, however, the interaction between SR Probability and Adjacency was also significant (*F*(1, 113) = 4.11, *p* = .045, *η^2^* = .003). Post-hoc comparisons (Holm-corrected) revealed that participants showed significantly higher ACC for High SR pairs compared to Low SR pairs in both the adjacent condition (*MD* = 0.03, *p* = .046) and the non-adjacent condition (*MD* = 0.07, *p* = .001). Regarding RTs, the ANOVA yielded a significant main effect of adjacency (*F*(1, 113) = 16.78, *p* < .001, *η^2^* = .004), indicating faster responses for adjacent pairs compared to non-adjacent pairs. However, neither the main effect of SR probability (*F*(1, 113) = 0.35, *p* = .558, *η^2^* < .001), nor the interaction between SR probability and adjacency (*F*(1, 113) = 1.09, *p* = .298, *η^2^* < .001) reached significance (**Supplementary Figure 2b,c**).

### 3.5. Cognitive flexibility and working memory predict successor representation in structural learning

Given established evidence that SR-based learning is cognitively demanding(Gershman, 2018), we hypothesized that the learning of SR-based syllable structure in the current study would depend on individual cognitive capabilities. To specifically quantify the effectiveness of SR learning while controlling for individual baseline differences, we calculated the differential ACC and RT between High SR Probability and Low SR Probability pairs across all pairs (including adjacent and non-adjacent). A higher positive Δ ACC (*Accuracy_Hig_*_ℎ_ *_SR_* -*Accuracy_Low SR_*) and more negative Δ RT (*Reaction Time_Hig_*_ℎ_ *_SR_* − *Reaction Time_Low SR_*), indicate stronger structural learning. We examine the relationship between SR learning effectiveness and cognitive flexibility as well as working memory.

Cognitive flexibility was measured using the Cognitive Flexibility Inventory (CFI, Dennis & Vander Wal, 2010), a self-report questionnaire evaluating the ability to perceive difficult situations as controllable and generate multiple alternative solutions. Participants were divided into a High-CFI group (n = 48) and a Low-CFI group (n = 47). Independent-samples t-tests with FDR correction revealed a significant difference in Δ ACC, with the High-CFI group (*M* = 0.21, *SD* = 0.13) performing better than the Low-CFI group (*M* = 0.12, *SD* = 0.18), *t*(83.2) = 2.80, *p_adj_* = .013, *Cohen’s d* = 0.580 (**Figure 6a**). No significant difference was observed in Δ RT between the groups, *t*(92.96) = 1.03, *p_adj_* = .303, *Cohen’s d* = 0.21 (**Figure 6b**).

**Figure 6.**
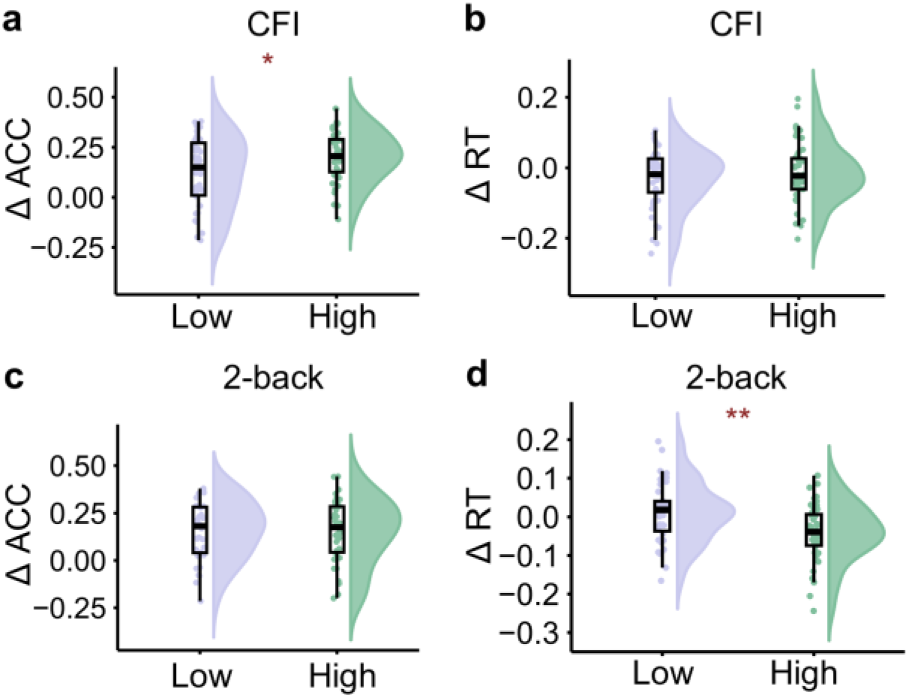
Impact of cognitive flexibility and working memory on SR performance (Δ ACC and Δ RT). The y-axes represent the magnitude of the SR effect, calculated as the difference in performance between High-SR and Low-SR pairs (i.e., Δ = High SR – Low SR). **(a)** Δ ACC by cognitive flexibility. High-CFI individuals showed significantly higher structural learning sensitivity (Δ ACC) than the Low-CFI group. **(b)** Δ RT by cognitive flexibility. No significant difference was observed. **(c)** Δ ACC by working memory. No significant difference was found. **(d)** Δ RT by working memory. The high working memory group exhibited a stronger SR effect (more negative Δ RT values) than the low group. Boxplots show the median (center line), the interquartile range (25^th^–75^th^ percentiles), and the minimum and maximum values within 1.5 × IQR. * *p* < .05, ** *p* < .01, *** *p* < .001.

Complementing the analysis of cognitive flexibility, working memory capacity was assessed using the 2-back task, a classic paradigm requiring participants to monitor a series of stimuli and respond when the current stimulus matches the one presented two steps earlier (Jaeggi et al., 2010; Kirchner, 1958). Participants were divided into High (n = 43) and Low (n = 43) working memory groups based on task performance. No significant difference was observed in Δ ACC between the groups, *t*(77.55)=0.21, *p_adj_* = .836, *Cohen’s d* = 0.05 (**Figure 6c**). However, a significant difference was found in Δ RT, with the High working memory group (*M* = −0.05, *SD* = 0.07) showing more negative Δ RT values than the Low group (*M* = 0.01, *SD* = 0.08)*, t*(81.67) = −3.35, *p_adj_* = .002, *Cohen’s d* = −0.720 (**Figure 6d**). Together, these results suggest that while cognitive flexibility is essential for the successful acquisition of SR syllable structures, SR learning is associated with higher cognitive capabilities: cognitive flexibility and working memory.

## 4. Discussions

The current study investigated the acquisition and the representational nature of the temporal structure embedded in syllables streams at a natural speech rate using an adapted auditory pyramid-structured paradigm (Tacikowski et al., 2024). Participants exhibited robust sequence learning, reliably distinguishing learned adjacent syllable pairs from the unlearned non-adjacent distractors. Moreover, we found a significant performance asymmetry: despite matched exposure frequencies, recognition accuracy was significantly lower for in-out pairs (low TP) than for out-in pairs (high TP), indicating sensitivity to transitional structure beyond simple associative pairing. Crucially, this structural knowledge generalized to non-adjacent syllable pairs that were never presented during learning, demonstrating flexible multi-step relational learning. Model-based analyses revealed that behavior performance was better captured by the transitional probabilities derived from a successor representation (SR) model than by a geometric (Euclidean) account. Finally, structure learning was modulated by individual cognitive capacity, higher cognitive flexibility and working memory associated with superior structure acquisition. Taken together, these findings suggest that humans rely on SR-like predictive representations to extract temporal structure from continuous speech at naturalistic rates, particularly for non-adjacent dependencies, extending cognitive map–based mechanisms from spatial navigation and visual sequence learning to the rapid auditory processing.

Our results showed that the learned adjacent pairs were recognized more accurately and faster than unlearned non-adjacent distractors, replicating prior findings (Ecker et al., 2013; Jones et al., 2023). Predictive processing may contributed to this facilitation, as the first syllable strongly predicted the subsequent syllable in learned adjacent pairs, potentially allowing pre-activation of expected representations and reducing processing uncertainty (Brothers et al., 2019). Such predictive facilitation is consistent with SR-based accounts, in which a multi-step transition structure supports efficient anticipation of upcoming elements. Conversely, correctly rejecting unlearned non-adjacent pairs may require a more cautious rejection strategy. Since the non-adjacent distractors shared constituent syllables with the learned pairs, they likely elicited familiarity-based interference that required additional cognitive control to resolve. Moreover, violations of predicted transitions may have generated prediction error signals, increasing decisional conflict and prolonging response latencies. Together, these factors may explain the lower accuracy and slower responses observed for non-adjacent pairs.

The significant performance asymmetry between adjacent pairs (in-out vs out-in) provides initial evidence for structural learning. We first interpreted this effect in terms of local, one-step transitional probabilities (TP) derived directly from the pyramidal topology. For any given connected pair (e.g., wo and mu), the phonetic properties and co-occurrence frequency are identical, whereas the directional predictability differs. Specifically, an outer syllable (e.g., wo) has only two permissible successors (e.g., mu, ka), yielding a TP of 0.50 for the wo-to-mu transition. In contrast, an inner syllable (e.g., mu) acts as a hub with four permissible successors (e.g., wo, ka, le, te), thereby diluting the TP for mu-to-wo to 0.25. Our observation that the recognition accuracy is driven by transitional probability aligns well with seminal work on statistical learning in continuous speech (Franco et al., 2015; Saffran et al., 1996). To extend this local, one-step account to multi-step transitions, we modeled TP with the successor representation (SR), which enables quantification of transition strengths for non-adjacent pairs that are undefined in standard Markovian models (Gershman, 2018). As shown in **Figure 5**, SR-derived TP robustly predicted recognition accuracy across both adjacent and non-adjacent conditions, demonstrating that a unified predictive mechanism underlies structural learning across multiple temporal scales. Critically, unlike earlier NAD studies using short and deterministic A-X-B frames (Endress & Bonatti, 2007; Wang et al., 2019), our design captures flexible, probabilistic, and multi-step structure that more closely approximates the complexity of natural continuous speech.

The TP-driven behavioral asymmetry we observed mirrors predictive biases reported in spatial navigation, where repeated unidirectional traversal skews hippocampal place fields toward upcoming locations (Mehta et al., 2000). These phenomena reflected the learning of SR probabilities. SR-based accounts propose that the hippocampal formation encodes a predictive map in which current states carry information about likely future states under the learned transition structure (Stachenfeld et al., 2017). These parallel raises the possibility that the hippocampus implements SR-like predictive computations that may also contribute to learning temporal structure in language. Human intracranial evidence further suggests a cortical–hippocampal hierarchy: early sensory regions primarily encode local transitional probabilities, higher-order cortical areas represent ordinal structure, and the hippocampus selectively integrates sequence identity and higher-order relational structure (Henin et al., 2021). Together, these findings raise the possibility that the hippocampus supports the construction and stabilization of predictive maps underlying syllable-sequence structure learning.

At the same time, such cross-domain generalization warrants caution. In our task, an explicit Euclidean (geometric) distance provided a less precise model accounting of behavior, whereas Euclidean structure is often salient and behaviorally relevant in spatial learning (e.g., metric coding and heading-direction representations) (Lv et al., 2024; Shine et al., 2016). More broadly, whether learners rely primarily on SR-like predictive maps, Euclidean geometry, or hybrid combinations likely depends on task demands and the representational affordances of the learning environment (Tacikowski et al., 2024). Accordingly, treating SR as a domain-general mechanism for sequence learning should be conditioned on context, rather than assuming a single representational format across domains.

Furthermore, we found that individuals with higher general cognitive ability, as indexed by working memory performance on the n-back task and scores on the cognitive flexibility questionnaire, acquired the sequential structure more effectively. This pattern aligns with prior evidence suggesting that the acquisition and expression of model-based or structure knowledge are constrained by cognitive resources (Hayes & Henderson, 2017; Lynn et al., 2020; Otto et al., 2015). Although the successor representation (SR) is often characterized as an intermediate form of representation between model-free and model-based learning, it nonetheless depend on control resources (Gershman, 2018). Supporting this view, Lynn et al. (2020) showed that a temperature-like parameter (β), derived from n-back task, shapes the predictive horizon of SR-like multistep map. Given that cognitive ability and executive functions are typically associated with the prefrontal cortex, while SR is linked to hippocampus, our findings motivate further investigation into how SR acquisition is supported by distinct prefrontal cortex and by prefrontal–hippocampal interactions, especially for speech processing. Note that the CFI measure reflects self-reported coping strategies rather than executive function in the strict behavioral sense. Future studies could therefore incorporate objective executive-function test (e.g., WCST/Stroop/Flanker) to more directly examine how executive flexibility contributes to SR-based structural learning.

## Limitation and future direction

An important open question concerns whether SR-like predictive representations in syllable-sequence structure require offline consolidation, and whether sleep selectively strengthens the multistep predictive horizon. Recent work further suggests that such SR-like predictive representations may be internally expressed and strengthened through replay dynamics during sleep and rest after learning (Kurth-Nelson et al., 2016). Future studies combining sequence learning paradigms with sleep or rest manipulations may clarify whether replay contributes to the strengthening and generalization of predictive representations.

Our findings also carry implications for early language acquisition. Young learners lack explicit linguistic knowledge to recognize NADs directly (Saffran, 2001; Saffran et al., 1996). To infer morphological rules and lexical structures embedded in words, they must rely on sensitivity to syllable-level transition regularities. Future work examining whether infants and children exhibit SR-consistent learning patterns may further elucidate the developmental foundations of speech segmentation and grammatical acquisition.

Several limitations of the present study should be acknowledged. First, inferences about representational format were derived from behavioral indices, which alone cannot uniquely determine the underlying representational code. Future neuroimaging approaches may help further distinguish the predictive, geometric, or hybrid representations. Second, although the structured tone sequences enabled controlled examination of statistical regularities, these stimuli are simplified relative to complex naturalistic inputs such as language or music, raising questions about ecological generalizability. Despite these constraints, the present findings demonstrate that humans can acquire and exploit structured relationships in rapidly unfolding syllable streams, providing a critical link between speech sequence learning and emerging theories of cognitive maps and predictive representations under ecologically relevant temporal demands.

## Author Contributions

Jiali Liu (Conceptualization, Funding acquisition, Supervision, Writing—original draft), Yixiang Wang (Data collection, Formal analysis), Jiayu Feng (Data collection, Formal analysis), Haichao Zhao (Supervision, Writing—review & editing), Qinghua He (Supervision, Writing—review & editing), Xuefeng Yang (Conceptualization, Supervision, Writing—review & editing), Baishen Liang (Conceptualization, Supervision, Writing—review & editing).

## Artificial intelligence

Artificial intelligence–assisted language editing tools were used to improve the clarity and readability of the manuscript, and polish the coding. No artificial intelligence tools were used in the research design, data collection, or interpretation of the results. Ethics: This study was approved by the Ethics Committee of Southwest University (Approval No. H2543). All procedures complied with the Declaration of Helsinki. All participants provided informed consent prior to participation.

## Preregistration

No aspects of the study were preregistered.

### Data, Materials, and Analysis Code Availability

All materials, anonymized data, and analysis scripts are available at the Open Science Framework: https://osf.io/y5dpq/?view_only=63841c46fb58441a84ade920f989dcc0

## Appendix

1. Successor representation transitional probability matrices derived from varying model parameters.

2. Reaction times for non-adjacent pairs by in-out, out-in, and out-out categories; Reaction times for adjacent and non-adjacent pairs with different successor representation transitional probabilities levels.

