## Supplementary materials for "Successor representation supports structural learning of syllable sequences"

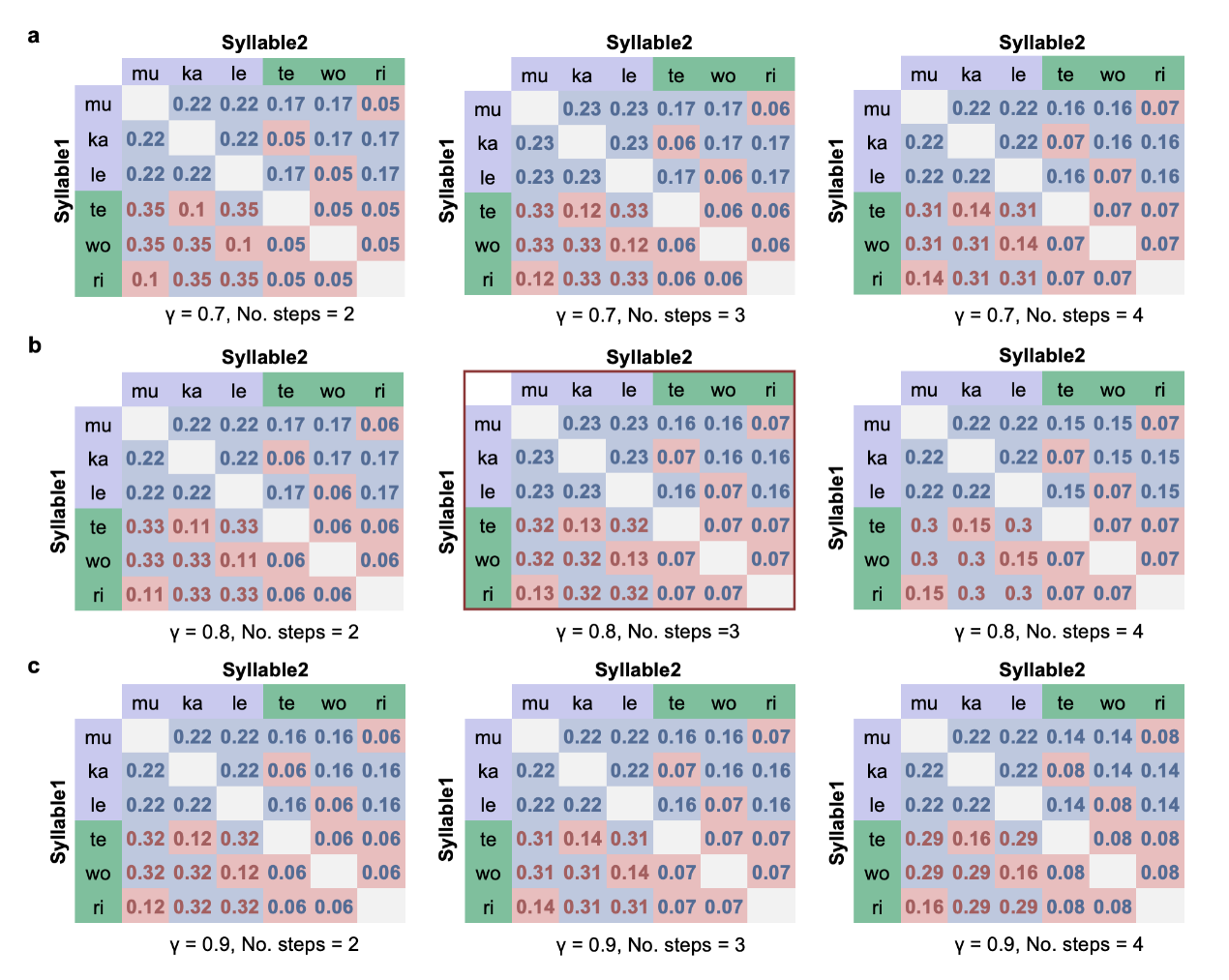

**Supplementary Fig. 1.** Successor representation (SR) transitional probability matrices derived from varying model parameters. The discount factor γ was set to 0.7 **(a)**, 0.8 **(b)**, and 0.9 **(c)**, while the steps varied across columns (Left: steps = 2, Middle: steps = 3, Right: steps = 4). Notably, the classification of High (red text) versus Low (grey-blue text) SR probability pairs remained consistent across all parameter combinations for both adjacent (gray shading) and non-adjacent (coral shading) conditions.

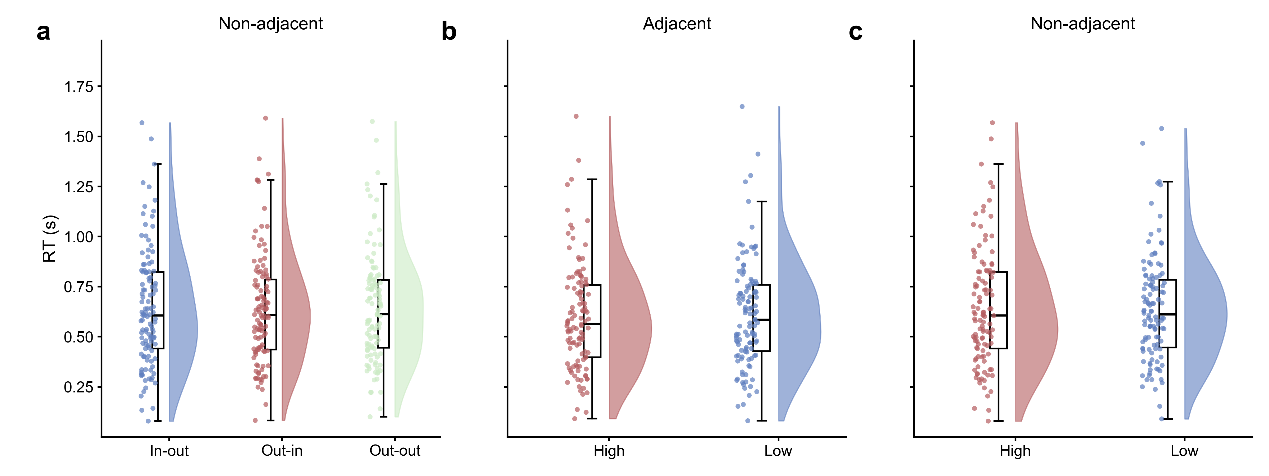

**Supplementary Fig. 2. (a)** Reaction times (RT) for non-adjacent pairs during the testing phase by in-out, out-in, and out-out categories; no significant differences were found across categories. **(b-c)** RT for trial pairs with different successor representation transition probability (TP) levels. For both adjacent **(b)** and non-adjacent **(c)** conditions, no differences were found between pairs with high and low TPs. Boxplots show the median (center line), the interquartile range (25^th^–75^th^ percentiles), and the minimum and maximum values within 1.5×IQR.
